# Pre-Breeding of Sorghum (Sorghum bicolor L. Moench) for Drought Tolerance in the Semi-Arid Zones of Nigeria

**DOI:** 10.64898/2026.08.10.743909

**Authors:** Muhammad Ahmad Yahaya, Mohammad Faguji Ishiyaku, Lateefat Bolanle Hassan

## Abstract

Genetic gains for yield and yield-contributing traits are low or stagnant in sorghum especially under low soil moisture environments which have contributed to a yield gap of 3.0 in Africa. Exploring the extent of variation for yield-determining traits in sorghum will effective variety design to boost production in drought-stressed environments. Therefore, the objective of this study was to determine genetic variability, heritability and genetic gains for agronomic and physiological traits in sorghum under varying drought stress conditions to guide cultivar and trait selection for breeding. The study was conducted as a three-way factorial treatment structure involving a genetically diverse panel of two hundred and twenty-five (225) sorghum genotypes which were grown under three drought conditions [i.e., non-stressed (NS), pre-anthesis drought stress (PrADS), and post-anthesis drought stress (PoADS)] and two environments (e.g., field and glasshouse environments) using a 15 x 15 alpha lattice design in two replicates. Data were collected on agronomic and physiological traits, namely: days to anthesis (DA), days to maturity (DM), plant height (PH), stay-green (SG), seed weight (TKW), biomass yield (BM), harvest Index (HI), grain yield (GY), leaf area (LA), leaf chlorophyll content (SPAD) and stomatal conductance (SC) and subjected to various statistical analyses. Combined analysis of variance showed highly significant (P < 0.001) genotype, drought condition and environment, and their interactions for most traits under assessment. Genotypic coefficient of variation (GCV) was lower than phenotypic coefficient of variation (PCV) for all traits. Heritability in the broad-sense (H^2^) was moderate for GY (40%) under PrADS condition, but high under NS (60%) and PoADS (65%) conditions. Similarly, high genetic advance was recorded for traits with high heritability. GY positively and significantly correlated with HI under NS (r = 0.88), PrADS (r = 0.62) and PoADS (r = 0.61). Other agronomic and physiological traits poorly correlated with GY. The following genotypes were selected based on high grain yield and suitable agronomic and physiological traits under NS conditions; DAN YARA (5.7 t/ha and SC = 335.5) and JARWA (GY = 5.6 t/ha, SC = 326.3); under PrADS CSRO1 (GY = 2.9 t/ha, SC = 275.2) and ICNSL2014-021-1 (GY = 2.7 t/ha, SC = 268.2) and under PoADS conditions DANYAR BANA (GY = 4.2 t/ha, SC = 237.2) and DAN YARA (3.9 t/ha. SC = 330.0). The selected genotypes with the desirable traits are useful genetic resources for breeding high-performing sorghum hybrids to boost sorghum productivity in drought-prone areas in Africa.

## INTRODUCTION

Sorghum (Sorghum bicolor [L.] Moench) is an important multi-purpose crop grown mostly as a cereal grain and animal feed. It serves as an important cereal for human diet attributed to its nutrient-rich grain with low fat content (3.3 %), high protein (11.3 %), fiber (6.3 %), vitamin B complex, and essential macro- and micronutrients (Ejeta and Knoll, 2007; Thilakarathna et al., 2022). In addition to its nutritive values, the crop residues are used as animal feed, and for producing fiber, biofuel, and, more recently, cellulosic ethanol feedstock (Assefa et al. 2010).

The continental Africa accounts the highest production of sorghum estimated at 28.2 million metric tons (FAOSTAT 2020). On the continent, sorghum is the primary cereal crop for small-scale farmers, particularly those living in the dry regions (Devnarain et al., 2016). The prevalent agricultural systems of the resource poor farmers rely wholly on rainfall as primary source of water. This reliance makes sorghum production vulnerable to the impacts of drought as the farmers lack essential irrigation systems (Devnarain et al., 2016). Therefore, drought has been reported to be the most important constraint to sorghum production in the dry regions of Sub-Saharan Africa (Assefa et al. 2010; Amelework et al., 2015; Yahaya and Shimelis, 2022).

Under pre-anthesis (panicle development) drought stress, where plants experience drought stress during panicle initiation before flowering, agronomic traits such as grain yield were reportedly reduced by 32 to 36%, 1000-seed weight by 23 to 31%, and plant height by 13 to 14% across different studies (Assefa et al., 2010; Emendack et al., 2014; Batista et al., 2019; de Souza 2020; Araki et al., 2022; Golzardi et al., 2023). Chlorophyll content in drought tolerant sorghum genotypes have been reported to be significantly higher than in drought sensitive genotypes under drought stress. In addition, stomatal closure and high leaf area index have been identified as adaptive trait to predict pre-flowering drought stress tolerance (Xu et al., 2010). Water stress in the post-anthesis stages (between flowering and grain development) have been reported to be more detrimental to sorghum crop yields and could reduce grain yield by 50 - 55% ( Assefa et al., 2010; Emendack et al., 2018; Batista et al., 2019). In addition, drought stress at this stage could reduce the leaf area by 11%, and plant height by 8.7% (Abraha et al., 2015). Genotypes resistant to both pre- and post-anthesis drought have been identified. However, only a few sorghum genotypes combine a high level of resistance of both types. Therefore, the adaptive drought-related traits should be exploited as a strategy in breeding grain sorghum cultivars for drought-prone environments.

It is necessary to have knowledge of genetic variability and heritability estimates in order to help incorporate these qualities into better grain sorghum cultivars or lines. Drought related traits in sorghum have been reported to be inherited and controlled by one or a few genes, while other traits are quantitatively inherited and controlled by many genes (Ludlow and Muchow, 1990). For example, earliness has been shown to be under genetic control and has high heritability. However, earliness has been reported to be frequently association with reduced yield. Narrow-sense heritability of plant height, stay-green and grain yield traits were found to be 44, 39 and 29% respectively for post-anthesis water stress (Mkhabela 1995). From the study, the estimates of narrow-sense heritability of the plant height and stay-green trait is sufficiently high to warrant a breeding program for post-flowering drought resistance based on the stay-green trait. In theory, selecting for characteristics that confer drought tolerance can be done concurrently with selecting for high grain yield. In drought-prone environment, selecting for certain morpho-phenological traits offers an advantage over directly selecting for yield if the trait’s heritability is larger than yield and the trait is associated with yield (Blum 1989). The selection procedure for traits which confer drought resistance can, in theory, be done as in selection for pest and disease resistance at the same time as selection for yield (Blum 1989). Selection for traits that confer resistance to drought under field condition is desirable because it incorporates the actual genotype, environment, and their interactions. Plant breeders would gain knowledge about selection and breeding methodology to be used in generating drought-resistant cultivars from genetic variation in pre- and post-anthesis drought resistance features.

The objective of the study was to assess the usefulness of morpho-physiological traits related to pre- and post-anthesis drought stress response as indirect selection indices of improved productivity under variable drought stress. The specific objectives were:

1. To determine genetic variation and estimate heritability of pre- and post-anthesis drought resistance traits in grain sorghum.
2. To investigate the interrelationship of pre- and post-anthesis drought resistance traits and grain yield in grain sorghum.

## MATERIAL AND METHOD

### Plant materials

Two hundred and twenty-five (225) sorghum genotypes assembled from diverse origins were used for the study. These comprised 235 landraces and 52 elite lines from the International Crops Research Institute for the Semi-arid Tropics-Kano (ICRISAT-KN), 15 registered cultivars, 22 elite lines and 83 landraces from the Institute for Agricultural Research (IAR), Samaru, Nigeria, nine elite lines from the African Centre for Crop Improvement (ACCI) in South Africa, and 21 genotypes from the United States Department of Agriculture, Agricultural Research Service, National Plant Germplasm System (USDA-ARS, NPGS). The names, codes, pedigree information, and genotype sources of origin are presented in **Supplementary Table S1**.

### Experimental sites

The experiment was conducted at the Samaru research station under the Institute for Agricultural Research (IAR), Ahmadu Bello University, (A.B.U) Zaria (Latitude 11^0^11’N and 7^0^38’E), Northern Guinea Savannah Zone in Nigeria.

### Experimental Design and Cultural Practices

The two hundred and twenty-five (225) sorghum genotypes were evaluated under non-stressed (NS), pre-anthesis drought stress (PreADS), and post-anthesis drought stress (PoADS) conditions under field and greenhouse environments using a 15 x 15 alpha lattice design in two replicates. The three water regimes and two environments resulted in six testing environments, namely: glasshouse and non-stressed (E1); glasshouse and pre-anthesis drought stress (E2); glasshouse and post-anthesis drought stress (E3); field and non-stressed (E4); field and pre-anthesis drought stress (E5); field and post-anthesis drought stress (E6). Planting was carried out during the off-season cropping season (October to March) in 2024/2025. The description of the field layout and agronomic management was described in Yahaya et al., (2023).

In the field, sorghum plants were planted in ridges on a two-row of 5 m plot, with 30 cm intra-row and 70 cm inter-row spacing. Each ridge was covered with rice straw mulch, and a surface drip irrigation system were installed down the center. Two seeds were planted and thinned to one plant two weeks after emergence. The fertilizer was applied at the following rates: 120 kg/ha of urea (18% N), 60 kg/ha superphosphate (6%, P_2_O_5_), and 60 kg/ha potassium chloride (12% K_2_O). To monitor soil moisture content under field conditions two tensiometer sensors (Decagon Ech10HS, Pullman WA USA) were inserted at two depth zones: above, at the active root zone, and below the root zone at depths of 250 mm and 500 mm, respectively. The sensors recorded field capacity and permanent wilting point values at 22% and 8% volumetric moisture content, respectively. Under the field conditions, pre- and post-anthesis drought stress was imposed, according to Reddy et al., (2009). PreADS was imposed at growth stage 3 (when about one-third of the total leaf area has fully developed) and continued to stage 6 (half bloom stage), at which half of the plants in the plot have flowered (Vanderlip, 1993). PoADS was imposed by withdrawing irrigation during the booting stage approximately 45 days after sowing (Vanderlip, 1993). In our case, PoADS was imposed between 95 and 105 days in the field. Weed control was performed manually, whereas sugarcane aphid [*Melanaphis sacchari* (Zehntner)] was controlled by spraying chlorpyrifos (Avima SA) at a recommended rate of 1 mL per 100 litres of water.

### Data collection

#### Agronomic Traits

Data on different traits were collected according to the standard methods stated in the IBPGR and ICRISAT (1993). Data on the days to anthesis (DA): the number of days from mean emergence date of the field to the date when 50% of plants in the field started flowering; plant height (PH): measure the average height of the row from base to the tip of the ear before harvest and after 50% flowering (cm); stay-green (SG): visual scoring using 1-5 scale where 1, as complete senescence and 5, as very slight senescent; 1000-seed weight (TKW): weight of 100 kernels at moisture content of 12% (g); biomass yield (BM): measured as fresh weight of the above ground plant part including the grain (t/ha); harvest Index (HI): the ratio of dried grain yield to the dried above ground biomass; grain yield (GY): grain yield will be measured on a per plot basis and converted into per hectare after adjusting to 12.5% grain moisture content (t/ha).

### Physiological Traits

Leaf area (LA): The leaf area estimates in the current study were made at flowering from fully expanded leaves. Data was obtained only from leaves from the upper half of the canopy. An allometric coefficient (0.772) was used to estimate leaf area (LA) from leaf size data collected during non-destructive measurements, according to the following formula Fracasso et al., (2016):

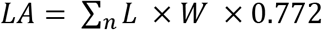

Where LA is leaf area, L is length, and W is the width.

### Leaf Chlorophyll Content

Leaf chlorophyll content was measured in actively growing leaves using a portable chlorophyll meter - The Soil-Plant Analyses Development (SPAD) (Model-502, Minolta Co., Ltd., Tokyo) as described by Dwyer et al. (1991). The SPAD values were taken at the middle of the leaf lamina of the second and fourth leaves from the top from three plants per plot (same plants that were used for stay green trait visual scoring), averaged on a plot basis for each leaf. SPAD values provide an indication of the relative amount of total chlorophyll present in plant leaves, based on the amount of light transmitted by the leaf (area 2 × 3 mm) in two wavelength regions in which the absorption of chlorophyll is different. Higher SPAD values represent higher total chlorophyll contents and the arbitrary SPAD values can be translated to the actual value of total chlorophyll per unit area (mg cm^−2^) using the equation.

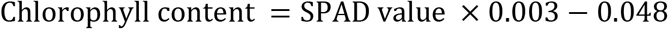

The leaf chlorophyll content measurements were taken when flag leaf emerged and measurements were repeated on the marked plants leaves at 2-week intervals until about 5 independent SPAD measurements were made (Xu et al. 2000).

### Leaf Stomatal Conductance

Stomatal conductance was measured with a leaf porometer (model SC-1, Decagon Devices Inc., Pullman, WA) on the abaxial surface of the mid portion of the youngest fully expanded leaf as mmol H2O m^-2^ s^-1^. For each sampled plant, the measurements of the leaf stomatal conductance were completed at two times at 7th and 21st days after anthesis with full clear air conditions at 10:00 am and 02.00 pm, in a sunny day. The relative humidity and air pressure at that time were 45% and 1000 mbar,

### Statistical Analysis

An analysis of variance (ANOVA) for all the recorded data and mean separation tests at the 5% and 1% levels of probability were performed using SAS software (version 9.2). The linear model of observations in an alpha lattice design was as follows:

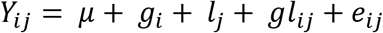

where *Y_ij_* is the observed mean yield for ith genotype at the jth location; *g_i_* is the genotype effect; *l_j_* is the effect of the location; *g_i_* × *l_j_* is the interaction effect of ith genotype at the jth location; and *e_ij_* is the residual error.

### Phenotypic and Genotypic Variance

Variability estimates including genotypic and phenotypic variances, heritability, genotypic and phenotypic coefficients of variations, and genetic advance were estimated according to (Lush 1949; Burton et al., 1953; Johnson et al., 1955).

These parameters were calculated according to the formula given by Lush (1949) for G x E Phenotypic Variance Component Model:

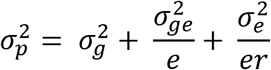

where 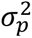 is the total phenotypic variance estimate, 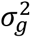 is the genotypic variance component,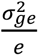 is GxE variance component, and 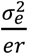 is the error variance component. The denominators *e* and *r* imply number of environments and replications, respectively.

The genetic parameters, mainly genotypic variance 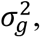, phenotypic variance 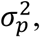, phenotypic coefficient of variation (PCV), and genotypic coefficient of variation (GCV), were derived by using the formula, suggested by Burton et al., (1953) and Johnson et al., (1955).

### Genotypic Variance Component

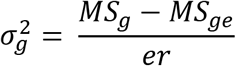

### Environmental Variance Component

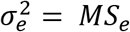

Genotypic coefficients of variation and phenotypic coefficients of variation were determined based on the method suggested by Burton et al., (1953) as follows:

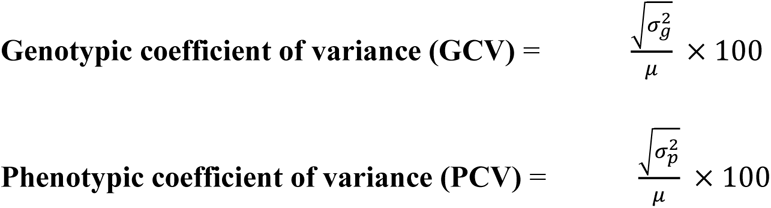

where *μ* is the grand mean of the trait, 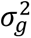 genotypic variance; 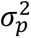 phenotypic variance; 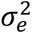 is environmental variance (error mean square from the ANOVA); MS*g* is the mean square of genotypes; MS*e* is the error mean square; and *r* is the number of replications.

Broad-sense heritability 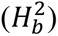 for traits across environments was estimated using variance components, according to Hallauer et al., (2010), as

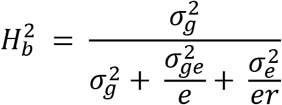

where 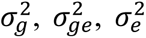 are the genotype, genotype x location, and residual variance components, respectively; *e* is the number of environments; and *r* is the number of replications.

The heritability estimate was categorized as described by Robinson et al., (1949) as follows:

0 - 30% = low, 30 - 60% = medium and >60% = high

### Estimation of Genetic Advance

The genetic advance at selection intensity (k) at 5% (2.06) was derived by using the following formula (Johnson et al., 1955):

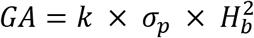

where GA represents the expected genetic advance under selection, *σ_p_* is the phenotypic standard deviation, 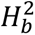 is heritability in broad sense, and k the selection intensity, or the value that is 2.06 at a 5% selection intensity.

### Estimation of Correlation Coefficients

Simple linear correlation coefficients (Pearson, 1895) were calculated to understand the relationship among the agronomic traits studied as below:

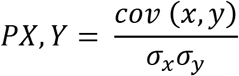

where *cov* is the covariance, *σ_x_* is the standard deviation of *x*, and *σ_y_* is the standard deviation of *y*. The corrplot package was used to run Pearson’s correlation coefficients in R version 4.2.0 (R Core Team, 2020).

## Results

### Effect of genotypes, drought conditions, and testing environments and their interaction on agronomic traits

The combined analysis of variance across locations showed highly significant (P < 0.001) genotype, environment, drought condition and their interaction effects for most of the measured traits (Table 1). The environment effect was non-significant (P > 0.05) for chlorophyll content (SPAD) and stomatal conductance (SC). Significant (P < 0.001) Genotype × environment interaction was recorded for most traits except days to anthesis (DA), days to maturity (DM), SPAD, SC and stay-green (SG). Genotype × environment × drought condition was significant for all measured traits except DA, and DM. The significant differences observed among the genotypes for most of the traits studied indicated the presence of large amount of genetic variability among the tested sorghum genotypes.

**Table 1:** Combined analysis of variance of grain yield and other agronomic traits among 225 sorghum genotypes evaluated under non-stressed (NS), pre-anthesis (PrADS) and post-anthesis (PoADS) drought stress conditions across two environments.

| Source of variation | Degree of freedom | Days to anthesis | Days to Maturity | Plant height (cm) | Leaf area (cm) | Chlorophyll content (mgcm<sup>-2</sup>) | Stomatal conductance (mmol m<sup>-2</sup> s<sup>-1</sup>) | Stay green |
| --- | --- | --- | --- | --- | --- | --- | --- | --- |
| Replication | 1 | 420.48 | 9847.96 | 2026.11 | 45235.76 | 2753.47 | 5927.81 | 0.34 |
| Environment (E) | 1 | 1070.58*** | 2204.13*** | 68892.31*** | 784388.89*** | 124.39ns | 497.07ns | 1.47* |
| Water regime (W) | 2 | 5336.66*** | 2212.91*** | 121170.00*** | 1631650.18*** | 10947.98*** | 100547.76*** | 233.57*** |
| Genotype (G) | 224 | 555.32*** | 1942.98*** | 40163.42*** | 106417.98*** | 1251.44*** | 55919.89*** | 9.76*** |
| G * W | 2 | 707.22*** | 2360.66*** | 13837.63*** | 135422.86*** | 4269.72*** | 28885.06*** | 2.00** |
| G * E | 224 | 5.16ns | 22.21ns | 1561.14*** | 10905.82*** | 29.16ns | 479.03ns | 0.08ns |
| G * E * W | 896 | 12.71ns | 36.31ns | 1989.49*** | 14714.57*** | 339.72*** | 2270.02*** | 0.99*** |
| Error | 1349 | 13.01 | 41.79 | 934.55 | 3616.84 | 38.67 | 431.09 | 0.32 |
| Mean |  | 65.43 | 117.82 | 225.08 | 350.30 | 41.57 | 179.81 | 2.59 |
| CV(%) |  | 5.51 | 5.49 | 13.58 | 17.17 | 14.96 | 11.55 | 22.04 |

| Source of variation | Degree of freedom | 1000- seed weight (g) | Biomass yield (t/ha) | Harvest Index | Grain weight (t/ha) |
| --- | --- | --- | --- | --- | --- |
| Replication | 1 | 14.11 | 180.56 | 0.09 | 0.42 |
| Environment (E) | 1 | 6721.11*** | 2537.11*** | 2.63*** | 65.45*** |
| Water regime (W) | 2 | 28347.38*** | 5052.54*** | 0.08*** | 856.18*** |
| Genotype (G) | 224 | 797.16*** | 24.83*** | 0.02*** | 7.11*** |
| G * W | 2 | 5190.66*** | 430.62*** | 0.04*** | 29.35*** |
| G * E | 224 | 149.26*** | 14.67*** | 0.01*** | 0.44*** |
| G * E * W | 896 | 72.71** | 11.10*** | 0.01*** | 0.66*** |
| Error | 1349 | 61.01 | 8.33 | 0.00 | 0.14 |
| Mean |  | 42.41 | 8.24 | 0.26 | 2.68 |
| CV(%) |  | 18.42 | 35.04 | 23.12 | 13.94 |

Agronomic and physiological responses among sorghum genotypes under non-stress (NS), pre-anthesis drought stress (PrADS) and post-anthesis drought stress (PoADS) in the greenhouse and field environments are presented in Table 2 and **Supplementary Table 2 (S2)**. Table 2 and Figure 1 showed the variability of DA in the studied sorghum genotypes across the two environments. Under NS condition, DA ranged from 42.5 for IS 8268 to 77.5 days for Wago Sane Red Sorghum. Genotypes IS 8268, ICNSL2014-022-8 and SSV20071012 recorded the lowest DA of ∼43, 49 and 51 days respectively. In contrast, Wago Sane Red Sorghum and Takumbo recorded the highest DA of 74 and 78 days under NS condition. Under PrADS condition, DA ranged from 46 for IS 8268 to 85 days for 12KNICSV-93. The genotypes 12KNICSV-93 and 12KNICSV-107-2 recorded the highest DA of 84 and 85 days under PrADS condition while genotypes IS 8268, ICNSL2014-022-8 and SSV20071012 recorded the lowest DA values across the environment. Under PoADS condition, DA range was from 43 for IS 8268 to 80.6 days for Takumbo. Genotypes IS 8268, ICNSL2014-022-8 and SSV20071012 recorded the lowest DA. Conversely, Takumbo and Yar’Fargore recorded the highest DA of 81 days under PoADS condition. The mean DA was 5 days earlier in NS condition (63 DAS) compared to PrADS condition (68 DAS), while it was one day earlier when compared to PoADS condition (64 DAS). Regardless of drought conditions, most of the tested sorghum genotypes recorded DA ∼50 to >70 days after planting (DAP) in both environments. The early maturing sorghum genotype under all drought conditions recorded the same DA regardless of environment. Overall, the genotypes took longer to reach anthesis under stress environments (PrADS and PoADS) than under NS conditions (Table S2 and Figure 1A).

**Figure 1:**
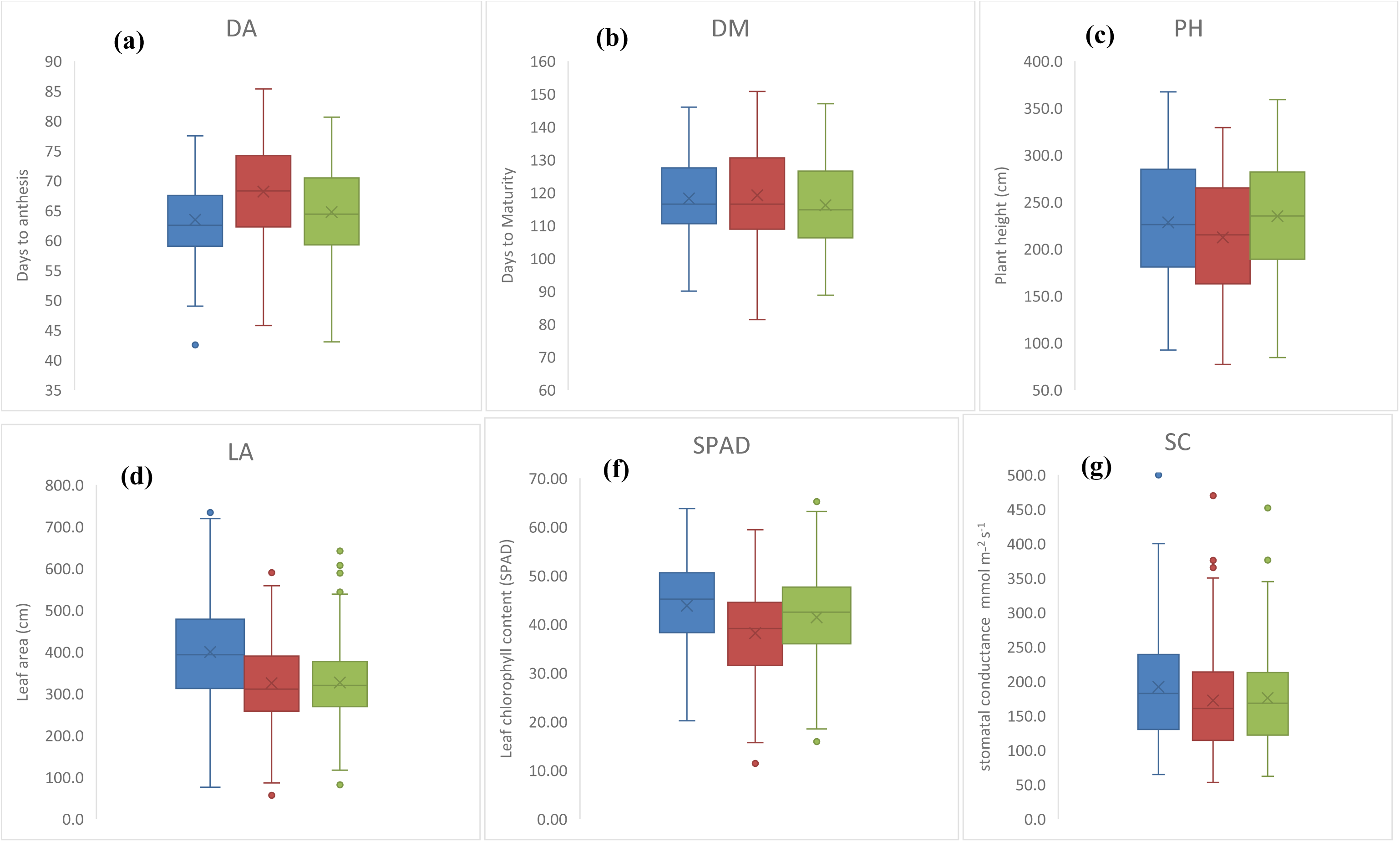
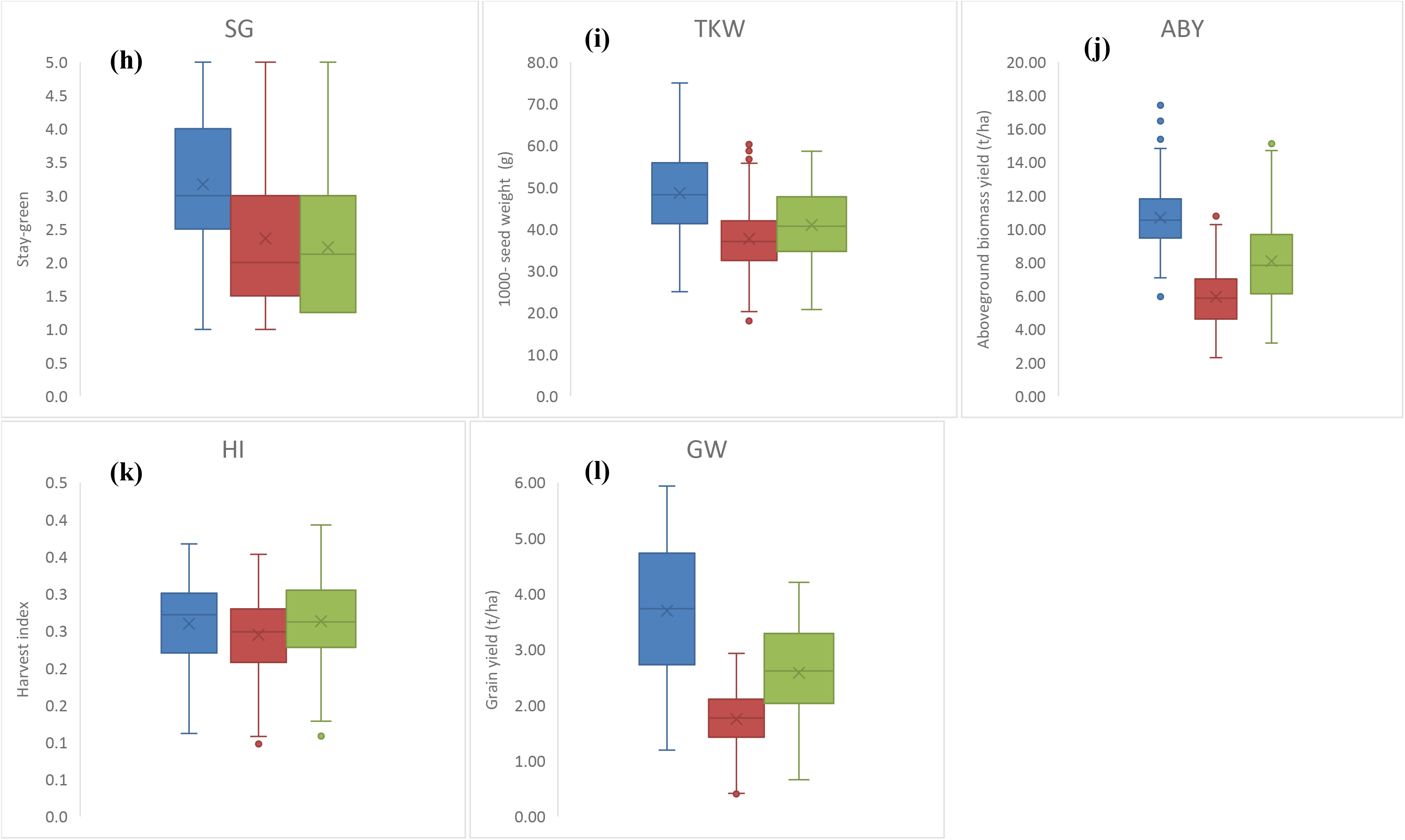
Box plots of the studied traits showing the mean performance of all the 225 sorghum genotypes under NS (Blue), PrADS (Orange) and PoADS (Ash) conditions.

**Table 2:** Mean performance of 225 sorghum genotypes evaluated under non-stressed (NS), pre-anthesis (PrADS) and post-anthesis (PoADS) drought stress conditions across two environments.

| Genotype | Days to anthesis |  |  | Days to maturity |  |  | Plant Height (cm) |  |  | Leaf area (cm) |  |  | Chlorophyll content (mgcm<sup>-2</sup>) |  |  |
| --- | --- | --- | --- | --- | --- | --- | --- | --- | --- | --- | --- | --- | --- | --- | --- |
|  | NS | PreADS | PoADS | NS | PreADS | PoADS | NS | PreADS | PoADS | NS | PreADS | PoADS | NS | PreADS | PoADS |
| S7-Lata/RIB/BC1-3-1-1-V | 65.00 | 78.33 | 65.50 | 115.00 | 116.38 | 112.50 | 286.75 | 253.75 | 223.13 | 433.33 | 500.64 | 606.94 | 36.57 | 36.45 | 44.84 |
| ICNSL2014-024-2 | 70.00 | 79.83 | 71.88 | 112.50 | 113.88 | 111.25 | 306.75 | 253.75 | 245.00 | 516.08 | 545.03 | 588.25 | 21.32 | 21.16 | 42.63 |
| GADAM | 60.00 | 55.83 | 57.13 | 110.00 | 111.38 | 102.50 | 136.50 | 146.00 | 149.88 | 324.47 | 298.52 | 403.87 | 43.46 | 42.22 | 47.46 |
| ICNSL2014-062 | 67.50 | 65.33 | 68.38 | 115.00 | 116.38 | 107.00 | 254.38 | 306.25 | 243.75 | 406.73 | 552.35 | 502.15 | 47.66 | 46.22 | 45.88 |
| CAPARLKSG20150308 | 61.25 | 71.83 | 71.50 | 120.00 | 121.38 | 112.50 | 166.00 | 167.50 | 231.00 | 132.95 | 344.82 | 501.41 | 53.51 | 51.77 | 47.64 |
| CAPARLGSG2015-0078 | 61.25 | 82.33 | 66.75 | 120.00 | 121.38 | 115.00 | 253.95 | 296.25 | 265.63 | 526.41 | 557.95 | 537.97 | 43.20 | 43.05 | 52.55 |
| ICNSL2014-025-8 | 66.25 | 79.83 | 72.00 | 109.50 | 110.88 | 101.25 | 296.38 | 284.25 | 268.75 | 531.02 | 398.97 | 341.88 | 49.92 | 48.36 | 37.76 |
| ICNSL2014-022-4 | 63.75 | 65.33 | 65.00 | 108.50 | 107.38 | 112.50 | 256.38 | 248.75 | 237.38 | 520.03 | 358.95 | 383.84 | 45.82 | 44.46 | 49.83 |
| YAR GUMEL | 60.00 | 74.08 | 62.50 | 115.00 | 116.38 | 112.50 | 267.63 | 229.75 | 202.25 | 452.08 | 370.11 | 479.61 | 55.86 | 54.02 | 50.41 |
| CF35:5 | 63.75 | 72.33 | 65.00 | 120.00 | 121.38 | 117.50 | 161.75 | 185.25 | 208.13 | 231.20 | 375.33 | 549.51 | 32.80 | 32.08 | 48.09 |
| ICSV111 | 60.00 | 61.08 | 59.38 | 111.00 | 112.38 | 106.25 | 188.25 | 193.75 | 189.88 | 268.70 | 475.62 | 353.78 | 45.10 | 43.78 | 38.54 |
| YARWASHA | 63.75 | 51.33 | 57.00 | 103.50 | 99.88 | 101.75 | 161.50 | 146.00 | 193.25 | 501.88 | 349.30 | 449.56 | 44.08 | 42.80 | 31.33 |
| ICSV 400 | 62.50 | 57.83 | 59.38 | 108.75 | 101.38 | 108.75 | 192.00 | 198.75 | 191.38 | 413.08 | 528.30 | 352.59 | 45.09 | 39.10 | 46.61 |
| ICNSL2014-065 | 66.25 | 57.83 | 61.25 | 90.00 | 88.88 | 113.75 | 241.25 | 290.00 | 249.25 | 532.17 | 592.80 | 472.29 | 49.29 | 53.95 | 52.93 |
| ICNSL2014-034 | 66.25 | 50.33 | 56.88 | 100.00 | 81.38 | 93.75 | 186.25 | 216.00 | 191.88 | 248.98 | 490.42 | 471.64 | 44.89 | 44.65 | 46.24 |
| CAPARLGSG2015-0035 | 60.00 | 75.33 | 65.50 | 113.75 | 113.88 | 111.25 | 232.50 | 285.50 | 257.88 | 442.94 | 509.68 | 351.09 | 53.61 | 51.87 | 49.29 |
| S7-Lata/RIB/BC1-1-17-1-V | 75.00 | 70.33 | 71.25 | 132.75 | 101.38 | 108.75 | 186.50 | 123.00 | 182.75 | 255.43 | 427.15 | 359.74 | 50.94 | 49.34 | 31.71 |
| AGUASASIN JAN'DAWA | 72.50 | 77.83 | 75.00 | 115.00 | 108.88 | 106.25 | 238.75 | 215.00 | 226.13 | 444.79 | 493.40 | 309.38 | 50.74 | 49.14 | 48.35 |
| MAI-RUWAN ZUMA | 71.25 | 67.83 | 69.38 | 115.00 | 106.38 | 102.50 | 303.75 | 316.25 | 303.75 | 465.69 | 420.99 | 461.51 | 51.76 | 50.12 | 43.00 |
| WAGO SANE RED SORGHUM | 77.50 | 77.83 | 76.88 | 131.25 | 108.88 | 102.25 | 276.25 | 326.25 | 283.63 | 548.08 | 341.14 | 391.64 | 52.89 | 51.19 | 43.57 |
| GAGARAWA - 3 | 73.75 | 70.33 | 72.50 | 119.75 | 101.38 | 98.75 | 313.75 | 329.00 | 292.00 | 492.64 | 506.07 | 343.85 | 46.23 | 44.85 | 38.84 |
| YAR'GETSO | 73.75 | 80.33 | 79.38 | 129.25 | 111.38 | 107.50 | 333.75 | 308.75 | 255.00 | 360.04 | 334.74 | 310.21 | 44.79 | 43.49 | 37.98 |
| YAR'FARGORE | 68.75 | 84.33 | 80.63 | 115.50 | 115.38 | 109.75 | 270.00 | 300.50 | 279.38 | 661.40 | 444.51 | 588.39 | 46.69 | 46.40 | 38.33 |
| ADAMAWA - 2 | 68.75 | 70.33 | 71.88 | 116.25 | 113.88 | 108.75 | 302.50 | 308.75 | 261.88 | 441.05 | 502.98 | 498.02 | 53.05 | 52.60 | 45.78 |
| WILD SORGHUM | 68.75 | 70.33 | 70.63 | 120.00 | 116.38 | 116.25 | 275.00 | 313.50 | 302.25 | 463.70 | 467.11 | 379.67 | 42.74 | 41.54 | 41.09 |
| MAI-GOJE | 67.50 | 80.33 | 74.50 | 115.00 | 116.38 | 128.88 | 257.50 | 199.50 | 227.50 | 491.33 | 550.40 | 350.10 | 43.26 | 42.02 | 32.64 |
| 12KNICSV-297-3 | 65.00 | 80.33 | 73.75 | 110.00 | 111.38 | 112.50 | 134.00 | 170.00 | 189.38 | 357.90 | 455.02 | 378.74 | 50.12 | 48.56 | 49.85 |
| 12KNICSV-297-1 | 62.50 | 70.33 | 65.75 | 113.75 | 111.38 | 110.00 | 159.00 | 153.00 | 180.63 | 463.33 | 351.84 | 376.13 | 40.37 | 40.30 | 47.61 |
| 12KNICSV-293 | 68.75 | 80.33 | 74.25 | 115.00 | 116.38 | 107.50 | 221.50 | 211.50 | 184.50 | 374.68 | 398.16 | 432.78 | 55.86 | 54.02 | 46.55 |
| 12KNICSV-297-2 | 75.00 | 80.33 | 77.25 | 117.50 | 116.38 | 112.50 | 197.00 | 103.75 | 99.13 | 339.28 | 509.79 | 417.86 | 50.43 | 48.85 | 42.54 |
| 12KNICSV-107-3 | 60.00 | 60.33 | 58.63 | 91.25 | 91.38 | 91.25 | 169.50 | 147.25 | 181.75 | 376.58 | 418.69 | 410.42 | 51.97 | 50.31 | 50.21 |
| 12KNICSV-295 | 62.50 | 65.33 | 63.75 | 95.00 | 96.38 | 92.50 | 139.00 | 127.50 | 152.13 | 559.43 | 428.08 | 396.64 | 51.07 | 50.70 | 40.30 |
| 12KNICSV-176 | 70.00 | 75.33 | 73.13 | 113.75 | 106.38 | 101.25 | 134.25 | 188.50 | 129.00 | 382.24 | 435.13 | 434.97 | 49.26 | 48.90 | 47.46 |
| 12KNICSV-297-4 | 68.75 | 80.33 | 77.50 | 115.00 | 111.38 | 105.00 | 136.25 | 122.00 | 217.63 | 536.08 | 295.01 | 468.66 | 58.12 | 56.16 | 51.27 |
| 12KNICSV-93 | 73.75 | 85.33 | 79.00 | 121.50 | 116.38 | 110.00 | 185.50 | 151.50 | 178.38 | 451.63 | 467.44 | 543.42 | 57.91 | 55.97 | 50.32 |
| 12KNICSV-260 | 72.50 | 70.33 | 70.63 | 118.75 | 111.38 | 108.75 | 231.75 | 201.25 | 154.00 | 384.08 | 305.83 | 360.81 | 50.33 | 48.75 | 53.38 |
| 12KNICSV-107-2 | 72.50 | 84.33 | 78.75 | 115.50 | 115.38 | 127.25 | 159.50 | 131.00 | 178.38 | 399.08 | 526.58 | 373.97 | 45.20 | 43.88 | 42.49 |
| 12KNICSV-418 | 65.00 | 70.33 | 68.75 | 113.75 | 111.38 | 107.50 | 167.50 | 143.25 | 148.75 | 382.80 | 531.53 | 486.37 | 53.66 | 53.25 | 47.73 |
| 12KNICSV-179 | 65.00 | 70.33 | 67.50 | 101.25 | 101.38 | 120.00 | 159.25 | 157.75 | 173.63 | 596.76 | 529.28 | 319.27 | 42.16 | 44.10 | 32.59 |
| CAPARLKSG20150293 | 67.50 | 75.33 | 71.00 | 105.00 | 106.38 | 101.25 | 219.63 | 109.00 | 231.25 | 632.50 | 492.19 | 455.63 | 57.13 | 56.65 | 52.21 |
| Environment (E) | *** | *** | *** | ns | *** | ns | ns | *** | *** | * | *** | *** | *** | *** | *** |
| Genotype (G) | *** | *** | *** | *** | *** | *** | *** | *** | *** | *** | *** | *** | *** | *** | *** |
| G x E | ns | ns | ns | * | ns | ns | ns | ns | *** | ns | *** | *** | *** | ns | ns |
| Mean | 63.43 | 68.14 | 64.71 | 118.18 | 119.17 | 116.10 | 228.29 | 212.21 | 234.73 | 399.46 | 325.04 | 326.39 | 43.80 | 38.18 | 41.40 |
| CV (%) | 6.22 | 4.71 | 5.60 | 5.11 | 5.71 | 5.62 | 7.92 | 11.38 | 18.54 | 12.07 | 20.33 | 19.61 | 13.15 | 14.34 | 17.08 |
| Tukey's HSD | 13.07 | 1.04 | 12.01 | 20.01 | 10.63 | 21.61 | 59.94 | 22.54 | 144.17 | 159.73 | 80.00 | 212.06 | 19.67 | 218.91 | 23.43 |

Table 2: Mean performance of 225 sorghum genotypes evaluated under non-stressed (NS), pre-anthesis (PrADS) and post-anthesis (PoADS) drought stress conditions across two environments (Continued).
| Genotype code | Stomatal conductance (mmol m<sup>-2</sup> s<sup>-1</sup>) |  |  | Stay green |  |  | 1000- seed weight (g) |  |  | Biomass yield (t/ha) |  |  | Harvest Index |  |  | Grain weight (t/ha) |  |  |
| --- | --- | --- | --- | --- | --- | --- | --- | --- | --- | --- | --- | --- | --- | --- | --- | --- | --- | --- |
|  | NS | PreADS | PoADS | NS | PreADS | PoADS | NS | PreADS | PoADS | NS | PreADS | PoADS | NS | PreADS | PoADS | NS | PreADS | PoADS |
| G1 | 232.0 | 221.5 | 214.3 | 3.0 | 4.0 | 2.6 | 36.5 | 31.0 | 31.0 | 8.8 | 10.8 | 6.8 | 0.3 | 0.2 | 0.3 | 3.7 | 2.3 | 2.8 |
| G2 | 390.7 | 372.5 | 253.5 | 4.0 | 3.0 | 3.0 | 37.5 | 33.8 | 34.6 | 12.3 | 4.5 | 8.3 | 0.3 | 0.2 | 0.3 | 4.8 | 1.5 | 3.1 |
| G3 | 290.4 | 277.1 | 177.3 | 2.0 | 2.0 | 2.8 | 55.5 | 42.0 | 39.8 | 9.3 | 3.6 | 3.8 | 0.1 | 0.2 | 0.2 | 1.2 | 0.6 | 0.8 |
| G4 | 253.7 | 242.2 | 205.9 | 3.0 | 4.0 | 2.6 | 57.0 | 50.8 | 49.0 | 11.5 | 6.8 | 6.8 | 0.2 | 0.2 | 0.2 | 2.6 | 1.2 | 1.8 |
| G5 | 382.7 | 364.9 | 276.6 | 1.0 | 2.0 | 2.1 | 47.0 | 40.8 | 39.4 | 12.0 | 3.6 | 9.3 | 0.3 | 0.3 | 0.2 | 4.6 | 1.4 | 2.2 |
| G6 | 141.8 | 135.7 | 213.0 | 3.0 | 4.0 | 2.6 | 59.5 | 49.8 | 46.8 | 13.3 | 7.4 | 7.4 | 0.3 | 0.3 | 0.3 | 4.6 | 2.3 | 2.4 |
| G7 | 204.3 | 195.2 | 246.5 | 4.0 | 3.0 | 2.4 | 69.5 | 60.3 | 57.4 | 10.5 | 7.6 | 7.4 | 0.3 | 0.2 | 0.3 | 4.4 | 2.0 | 2.5 |
| G8 | 134.8 | 129.1 | 198.4 | 4.0 | 5.0 | 3.5 | 62.0 | 52.3 | 48.8 | 11.1 | 4.9 | 7.4 | 0.3 | 0.4 | 0.4 | 5.0 | 2.7 | 3.8 |
| G9 | 296.5 | 278.9 | 249.9 | 5.0 | 4.0 | 3.3 | 57.0 | 51.0 | 50.9 | 12.4 | 8.8 | 5.8 | 0.3 | 0.2 | 0.3 | 5.9 | 2.0 | 2.4 |
| G10 | 233.4 | 219.7 | 196.4 | 1.0 | 1.0 | 3.3 | 51.5 | 47.3 | 47.8 | 10.4 | 4.1 | 5.6 | 0.3 | 0.3 | 0.3 | 5.2 | 1.6 | 2.7 |
| G11 | 184.2 | 173.6 | 133.6 | 2.0 | 3.0 | 1.8 | 69.5 | 41.0 | 34.7 | 12.3 | 6.6 | 5.3 | 0.3 | 0.3 | 0.3 | 4.4 | 2.1 | 2.3 |
| G12 | 118.4 | 111.8 | 147.9 | 3.0 | 3.0 | 3.0 | 50.5 | 47.0 | 48.3 | 10.6 | 5.1 | 6.5 | 0.2 | 0.2 | 0.2 | 2.7 | 1.2 | 1.3 |
| G13 | 176.0 | 165.8 | 181.1 | 4.0 | 4.0 | 2.6 | 52.5 | 36.2 | 35.6 | 11.6 | 6.0 | 6.0 | 0.3 | 0.3 | 0.3 | 4.8 | 2.0 | 2.7 |
| G14 | 207.6 | 195.5 | 205.1 | 4.0 | 4.0 | 3.3 | 68.0 | 54.8 | 48.8 | 12.5 | 7.9 | 11.7 | 0.2 | 0.2 | 0.2 | 3.8 | 1.6 | 2.4 |
| G15 | 297.0 | 283.4 | 226.5 | 1.0 | 3.0 | 3.0 | 63.0 | 53.8 | 50.6 | 14.7 | 5.1 | 9.2 | 0.3 | 0.3 | 0.3 | 5.4 | 1.9 | 3.3 |
| G16 | 166.1 | 156.6 | 193.6 | 3.0 | 3.0 | 3.0 | 59.5 | 39.8 | 37.4 | 13.5 | 6.4 | 5.4 | 0.3 | 0.2 | 0.3 | 4.8 | 1.5 | 2.8 |
| G17 | 223.2 | 210.1 | 186.1 | 1.0 | 3.0 | 3.0 | 68.5 | 37.5 | 32.0 | 13.6 | 9.9 | 5.7 | 0.2 | 0.1 | 0.3 | 3.3 | 1.4 | 1.7 |
| G18 | 167.7 | 158.0 | 238.0 | 4.0 | 4.0 | 3.3 | 54.3 | 52.3 | 51.9 | 12.6 | 5.9 | 9.1 | 0.3 | 0.3 | 0.2 | 5.0 | 1.9 | 2.0 |
| G19 | 154.9 | 146.1 | 138.0 | 3.0 | 3.0 | 3.0 | 50.0 | 32.5 | 35.0 | 12.5 | 7.7 | 5.4 | 0.2 | 0.2 | 0.2 | 2.8 | 1.4 | 1.2 |
| G20 | 107.1 | 101.3 | 94.4 | 4.0 | 4.0 | 3.3 | 58.0 | 51.5 | 51.0 | 11.6 | 9.1 | 8.6 | 0.3 | 0.2 | 0.2 | 3.9 | 1.6 | 2.1 |
| G21 | 216.3 | 203.7 | 146.7 | 4.0 | 4.0 | 2.6 | 40.0 | 36.8 | 38.9 | 11.7 | 7.0 | 4.3 | 0.2 | 0.2 | 0.2 | 2.5 | 1.2 | 1.0 |
| G22 | 176.7 | 169.0 | 120.5 | 3.0 | 3.0 | 3.8 | 59.0 | 52.0 | 51.5 | 10.4 | 6.3 | 5.9 | 0.2 | 0.2 | 0.2 | 2.5 | 1.3 | 1.9 |
| G23 | 148.4 | 142.1 | 281.9 | 4.0 | 4.0 | 3.3 | 36.5 | 27.0 | 30.6 | 13.5 | 8.1 | 10.8 | 0.2 | 0.2 | 0.2 | 4.1 | 1.3 | 2.0 |
| G24 | 94.6 | 90.9 | 274.9 | 3.0 | 3.0 | 2.4 | 48.0 | 39.0 | 37.5 | 10.5 | 8.0 | 6.1 | 0.2 | 0.2 | 0.2 | 2.2 | 1.3 | 1.0 |
| G25 | 121.4 | 116.3 | 186.4 | 4.0 | 3.0 | 2.4 | 52.0 | 51.5 | 50.3 | 11.8 | 5.9 | 8.4 | 0.2 | 0.3 | 0.2 | 3.2 | 2.4 | 2.6 |
| G26 | 196.2 | 187.5 | 304.3 | 4.0 | 5.0 | 3.5 | 58.5 | 52.5 | 51.9 | 12.4 | 7.5 | 10.3 | 0.2 | 0.1 | 0.1 | 2.8 | 0.9 | 1.4 |
| G27 | 128.7 | 123.3 | 294.3 | 5.0 | 4.0 | 4.0 | 40.0 | 27.5 | 32.7 | 11.1 | 4.6 | 5.1 | 0.3 | 0.2 | 0.3 | 3.7 | 0.9 | 2.2 |
| G28 | 145.2 | 139.0 | 166.2 | 4.0 | 4.0 | 3.3 | 66.0 | 40.8 | 39.8 | 11.9 | 4.7 | 7.7 | 0.3 | 0.2 | 0.3 | 5.2 | 1.3 | 2.5 |
| G29 | 78.6 | 75.7 | 136.2 | 5.0 | 5.0 | 4.3 | 49.5 | 34.8 | 35.0 | 11.4 | 3.2 | 4.9 | 0.1 | 0.2 | 0.1 | 1.5 | 0.6 | 0.7 |
| G30 | 146.3 | 140.0 | 100.1 | 4.0 | 5.0 | 4.3 | 61.5 | 38.3 | 32.8 | 11.1 | 2.3 | 4.0 | 0.2 | 0.2 | 0.2 | 2.5 | 0.4 | 1.0 |
| G31 | 131.2 | 125.7 | 218.5 | 5.0 | 5.0 | 4.3 | 55.0 | 48.5 | 48.0 | 10.1 | 3.4 | 6.4 | 0.3 | 0.3 | 0.3 | 3.5 | 1.2 | 2.3 |
| G32 | 187.4 | 179.1 | 271.7 | 5.0 | 5.0 | 5.0 | 41.0 | 29.3 | 29.6 | 11.4 | 3.9 | 4.6 | 0.3 | 0.3 | 0.4 | 4.1 | 1.7 | 2.5 |
| G33 | 281.3 | 268.4 | 196.3 | 4.0 | 5.0 | 4.3 | 31.0 | 20.2 | 25.7 | 10.6 | 3.1 | 5.6 | 0.1 | 0.2 | 0.2 | 1.6 | 0.8 | 1.0 |
| G34 | 297.4 | 283.7 | 197.2 | 4.0 | 4.0 | 4.0 | 50.0 | 44.8 | 44.6 | 10.6 | 3.0 | 6.0 | 0.1 | 0.2 | 0.2 | 1.5 | 0.9 | 0.9 |
| G35 | 148.3 | 142.0 | 119.6 | 5.0 | 4.0 | 4.0 | 58.5 | 39.8 | 43.3 | 12.8 | 2.8 | 7.7 | 0.2 | 0.3 | 0.2 | 3.3 | 1.0 | 1.3 |
| G36 | 280.1 | 267.3 | 182.8 | 5.0 | 5.0 | 5.0 | 47.5 | 39.8 | 37.3 | 9.7 | 3.0 | 6.1 | 0.2 | 0.4 | 0.2 | 2.4 | 1.6 | 1.8 |
| G37 | 189.3 | 181.0 | 108.6 | 5.0 | 5.0 | 4.3 | 49.0 | 41.3 | 39.5 | 9.2 | 6.6 | 6.0 | 0.3 | 0.2 | 0.3 | 3.5 | 1.8 | 1.9 |
| G38 | 211.0 | 198.7 | 162.1 | 5.0 | 5.0 | 5.0 | 65.0 | 56.8 | 54.5 | 9.5 | 3.8 | 5.1 | 0.3 | 0.3 | 0.3 | 4.1 | 1.8 | 2.7 |
| G39 | 142.3 | 134.3 | 194.9 | 5.0 | 5.0 | 2.9 | 37.5 | 34.8 | 36.3 | 13.9 | 3.7 | 5.5 | 0.3 | 0.3 | 0.3 | 5.0 | 1.9 | 2.8 |
| G40 | 228.1 | 214.7 | 309.6 | 5.0 | 5.0 | 3.5 | 48.0 | 40.8 | 39.9 | 11.2 | 4.0 | 9.4 | 0.3 | 0.3 | 0.2 | 4.6 | 2.1 | 2.7 |
| Environment (E) | *** | *** | *** | ns | ns | *** | * | *** | ns | *** | *** | *** | *** | *** | *** | *** | ns | *** |
| Genotype (G) | *** | *** | *** | *** | *** | *** | *** | *** | *** | *** | *** | *** | *** | *** | *** | *** | *** | *** |
| G x E | ns | *** | ns | ns | ns | ns | *** | ** | ns | ns | ns | *** | * | ns | *** | *** | * | *** |
| Mean | 191.8 | 171.9 | 175.7 | 3.2 | 2.4 | 2.2 | 48.6 | 37.7 | 40.9 | 10.7 | 6.0 | 8.1 | 0.3 | 0.2 | 0.3 | 3.7 | 1.8 | 2.6 |
| CV (%) | 2.9 | 3.4 | 19.6 | 15.3 | 23.3 | 26.9 | 12.3 | 23.2 | 20.5 | 29.1 | 42.4 | 36.8 | 20.4 | 25.2 | 23.7 | 12.8 | 17.9 | 11.8 |
| Tukey's HSD | 18.4 | 18.1 | 113.8 | 1.6 | 19.3 | 2.0 | 19.8 | 1.8 | 27.8 | 10.3 | 29.0 | 9.9 | 0.2 | 8.6 | 0.2 | 1.6 | 1.0 | 1.0 |

Data presented in Table 2 indicated that, the effect of water regime and genotypes had significant differences in DM. Under NS, The DM data ranged from 90 for ICNSL2014-065 to 146 days for Buk Wakana, while the range under PrADS condition was 81 days for ICNSL2014-034 to 151 days for Buk Wakana PrADS conditions. The lowest DM (∼ 90 days) was observed for ICNSL2014-065 and 12KNICSV-107-3 under NS and PrADS conditions, compared with Buk Wakana, Mai Bako Kono and Harjiu which were late maturing and recorded ≥140 days to maturity under NS and PrADS conditions. Under PoADS, days to maturity ranged from 89 days for E 29 to 147 days for Buk Wakana (Table 2 and Figure 1). Under PoADS condition, E 29, 12KNICSV-107-3 and 12KNICSV-295 were early maturing (∼93 days), compared with Buk Wakana, Harjiu, Mai Bako Kono which was late maturing (145 days) (Table S2 and Figure 1B).

For PH, data under NS condition ranged from 92.3 for AS 152 to 387.3 cm for Ndu Vari. The genotypes AS 152 (92.3 cm), P9402 (100.8 cm), CAPARLGSG20150111-1 (102.8 cm), GTPP7R(H)C5 (103.8 cm) and GP11BR (108.8 cm) recorded the lowest PH values whereas, NDU VARI (367.3 cm), YALAI (341.8 cm) and KITSE KAZA (336.3 cm) recorded the highest values (Table 2 and Figure 1). Under PrADS condition the values ranged from 77 for AS 71 to 329 cm for Gagarawa – 3. The lowest values for PH under PrADS conditions were recorded for AS 71 (77.0 cm), GTPP7R(H)C5 (83.2 cm), P9402 (85.6 cm), and CAPARLGSG20150111-1 (102.8 cm) whereas the genotypes GAGARAWA - 3 (329.0 cm), WAGO SANE RED SORGHUM (326.3 cm), and CAPARLGSG2015-0055 (325.0 cm) recorded the highest PH values under PrADS conditions. Under PoADS condition, AS 152 was the shortest (84.3 cm), whereas the highest PH was recorded by Bassa Dawa (359.0 cm) (Table 2). The lowest value for PH under PoADS condition was recorded for AS 152 (84.3 cm), 12KNICSV-297-2 (99.1 cm), E 29 (101.3 cm), and P9402 (107. 3 cm) whereas genotypes BASSA DAWA (359.0 cm), TSAWAN ZAKARA (355.9 cm), and KAFI MORI (352.2 cm) recorded the highest PH value (Table S2 and Figure 1C).

The LA differed significantly among studied genotypes under all studied drought conditions. The lowest LA was recorded for Jawar (75.7 cm), Juar (76.9 cm), Wiley (92.3 cm) and GPP4BR(H)C5 (100.9 cm) whereas the genotypes, SAMSORG 6 (733.5 cm), Fara Fara Kyal-Kyal (718.8 cm), and SAMSORG 14 (718.2 cm) recorded the highest LA value under NS condition. The LA values ranged from 56.1 for Jawar to 592.8 cm for ICNSL2014-065 under PrADS conditions. the genotypes with lowest LA value include JAWAR (56.1 cm), Juar (85.8 cm) and Wiley (114.5 cm) whereas ICNSL2014-065 (592.8 cm), CSRO1 (589. 7 cm) and CAPARLGSG2015-0078 (557.9 cm) recorded the highest values under PrADS conditions. Under PoADS conditions, LA values ranged from 81.2 for JAWAR to 641.5 cm for Sambulmu-3. Highest values were recorded for Sambulmu-3 (641. 5 cm), S7-Lata/RIB/BC1-3-1-1-V (606.9 cm), and Yar’Fargore (588,4 cm) and lowest values were recorded by the following genotypes Jawar (81.2 cm), Wiley (116.7 cm), and E 29 (116.9 cm) (Table S2 and Figure 1d).

Significant differences (P<0.05) in leaf chlorophyll content (SPAD) were recorded among the genotypes (Table 4). The SPAD values of the genotypes differed between the three water regimes, and PrADS condition significantly reduced the SPAD of all the genotypes (Mean = 38.2 mgcm^-2^). Under NS condition, Kaura Yellow Glume (63.8 mgcm^-2^), E 41 (61.4 mgcm^-2^) and AS 71 (59.9 mgcm^-2^) maintained the highest SPAD; while P9402 (20.2 mgcm^-2^), ICNSL2014-024-2 (21.3 mgcm^-2^), and SAMSORG 3 (21.9 mgcm^-2^) had the lowest values of this parameter (Table 2). Among the genotypes studied, the SPAD of the leaves ranged from 11.4 for E 29 to 59.4 mgcm^-2^ for AS 71 under PrADS condition. Lowest SPAD values were recorded by the genotypes E 29 (11.4 mgcm^-2^), Ako Variety (11.5 mgcm^-2^), and P9402 (15.7 mgcm^-2^) whereas highest values were recorded for AS 71 (59.4 mgcm^-2^), CAPARLKSG20150293 (56.7 mgcm^-2^), and 12KNICSV-297-4 (56.2 mgcm^-2^) under PrADS condition. Under the PoADS condition, genotype Ako Variety (15.9 mgcm^-2^), Farin Illo (18.5 mgcm^-2^), and P9402 (19.4 mgcm^-2^) had lowest SPAD compared to E 119 (66.0 mgcm^-2^), Kaura Yellow Glume (65.2 mgcm^-2^), and PATO (65.2 mgcm^-2^) which recorded the highest value (Table S2 and Figure 1e).

A significant variation (P<0.05) of SC was observed among the tested sorghum genotypes under the three water regimes (Table 2). The SC was significantly lower under water stressed treatments (PrADS and PoADS) than under NS condition and showed a general decreasing pattern with drought stress intensity (Table S2). Under the NS condition, SAMSORG 7 (499.8 mmol m-^2^ s^-1^) had the highest SC; followed by AS 152 (399.9 mmol m-^2^ s^-1^) and ICNSL2014-024-2 (390.7 mmol m-^2^ s^-1^). The lowest SC was recorded by IS 8265 under all water regimes. On the other hand, the maximum SC under PrADS condition was observed in SAMSORG 7 (469.5 mmol m-^2^ s^-1^), followed by AS 152 (375.8 mmol m-^2^ s^-1^) (Table 4). Under the PoADS condition, SAMSORG 7 (451.7 mmol m-^2^ s^-1^) recorded the highest SC followed by Mori Masaba (375.8 mmol m-^2^ s^-1^) (Table 2 and Figure 1f).

The stay green (SG) trait exhibited significant differences under all drought conditions among the tested sorghum genotypes (Table 4). Under NS conditions, 12% (28 genotypes) recorded the highest SG value (5.0) while about 1% (10 genotypes) recorded SG value of ∼1.0 (Table 4). Under PrADS and PoADS conditions, 6.7% (15 genotypes) and 1.3% (3 genotypes) recorded the highest SG value (5.0) while ∼8.7% (42 genotypes) recorded lowest SG value (1.0) under PrADS condition and ∼28.4% (64 genotypes) recorded lowest values of 1.25 (Table 2 and Figure 1g).

There was a significant difference in TKW between sorghum genotypes under NS and drought stress conditions. Data obtained ranged between 25 g (Juar) to 75 g (SAMSORG 44) for NS plants. Lowest TKW values were recorded for Juar (25.0 g), followed by Bog Farwa (29.5 g) and JAWAR (29.5 g) whereas genotypes, SAMSORG 44 (75.0 g), SAMSORG 46 (71.5 g), ICSV111 (69.5 g) and ICNSL2014-025-8 (69.5 g) recorded the highest values for TKW under NS conditions. TKW values for PrADS ranged from 18 G (E 29) to 60.8 g (SAMSORG 14), and the lowest values were recorded by E 29 (18.0 g), 12KNICSV-176 (20.2 g), and ICNL2014 026-8 (20.5 g) and the highest by SAMSORG 14 (60.8 g), ICNSL2014-025-8 (60.3 g) and SAMSORG 6 (60.0 g) (Table S2 and Figure 1h). Under PoADS, the TKW values ranged from 20.8 g (E 29) to 58.6 g (SAMSORG 14). Under NS condition, BM data ranged from 5.95 t/ha for SAMSORG 17 to 17.4 t/ha for Tun Buman Maiduguri, and under PrADS condition the values ranged from 2.3 t/ha for 12KNICSV-297-2 to 10.8 t/ha for S7-Lata/RIB/BC1-3-1-1-V. Under PoADS condition, BM value ranged from 3.2 t/ha for AS 97 to 15.1 t/ha for CSRO2 (Table 4). Lowest values under TKW were recorded for E 29 (20.8 g), followed by ICNL2014 026-8 (21.0 g) and SAMSORG 3 (22.1 g) whereas highest values were recorded for SAMSORG 14 (58.6), Sambulmu-3 (58.4) and Jar Balakwama (58.1 g).

Table 2 revealed considerable difference in grain yield (GY) among the tested sorghum genotypes across locations. The NS condition showed the best performance in GY with a mean of (3.7 t/ha) followed by PoADS condition (mean = 2.6 t/ha) and the least performance was shown by the PrADS condition (mean = 1.8 t/ha). GY under NS ranged from 1.2 for Gadam to 5.9 t/ha for Kaura Short Panicle-1, and 51% (114) of the tested sorghum genotypes had a higher GY (> 3.7 t/ha) and above the mean value. The genotype Kaura Short Panicle-1 (5.9 t/ha) produced maximum yield followed by Yar Gumel (5.9 t/ha), Dan Yara (5.7 t/ha) and ICNSL2014-023-5 (5.6 t/ha). Lowest yield under NS was produced by Gadam (1.2 t/ha), Kaura - 1 (1.3 t/ha), Yar Koma (1.3 t/ha) and KAURA - 2 (1.3 t/ha). Under PrADS, GY ranged from 0.4 for 12KNICSV-297-2 to 2.9 t/ha for CSRO1 and PrADS, 52% (116) of the tested genotypes recorded GY above the mean (1.75 t/ha). Under PrADS condition, the genotypes CSRO1 (2.9 t/ha) produced the highest yield followed by Tun Buman Maiduguri (2.8 t/ha), ICNSL2014-021-1 (2.7 t/ha) and Gwaza Banji Borno (2.7 t/ha). Lowest yield was recorded by 12KNICSV-297-2 (0.4 t/ha), AS 97 (0.4 t/ha) and GADAM (0.6 t/ha). Under PoADS condition, GY ranged from 0.7 for 12KNICSV-293 to 4.2 t/ha for Danyar Bana (Table 3). Across location, under PoADS, 51.1% (115) tested genotypes recorded grain yield performance above the mean value (2.58 t/ha). The highest GY value was recorded for Danyar Bana (4.2 t/ha), GAGARAU - 4 (4.2 t/ha), Kaura Short Panicle-1 (4.1 t/ha) and Kaura Mai Baki Kona (4.0 t/ha) whereas the genotypes 12KNICSV-293 (0.7 t/ha), Gadam (0.8 t/ha), and AS 97 (0.9 t/ha) recorded the lowest GY under PoADS condition (Table 2 and Figure 1i).

The HI data differed significantly (P < 0.001) among studied sorghum genotypes across drought conditions (Table 4). Under NS condition, genotypes Ndu Vari (0.11) recorded the lowest HI values, whereas ICNSL2014-023-5 (0.37) recorded the highest values. Genotype ICNSL2014-023-5 recorded the highest HI of 0.35 compared with the lowest value of 0,10 recorded for SAMSORG 42 under PrADS condition. Under PoADS condition, Kaura – 2 recorded the lowest HI values (0.11), whereas P9402 recorded the highest HI (0.39) (Table 4).

### Genetic Parameters

The variations among the tested genotypes for the target traits allowed for the selection of desirable genotypes for future crop improvement. In the current study, the investigation of covariance (environmental, genotypic and phenotypic), the genotypic- and phenotypic coefficient of variation (GCV and PCV), broad sense heritability and response to the selection for eleven recorded traits were shown in Table 5. From the results, under all conditions (NS, PrADS, and PoADS) the trait HI exhibited the lowest environmental variance 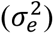 of 0.0 whereas LA recorded the highest value (2324.2, 4365.2 and 4096.5 respectively). Under all conditions, the phenotypic variance 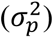 varied from 0.0 for HI to 6327.0 (NS), 3188.4 (PrADS) and 1819.3 (PoADS) for LA. Under NS condition, GxE interaction variance 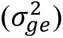 varied from -472.3 for leaf area to 8.7 for SPAD, whereas under PrADS condition, the least value was recorded for PH (-51.9) and topmost value was recorded for LA (622.2). Under PoADS conditions, the 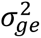 varied -7.0 for SPAD to 2174.3 for LA. Under all conditions, the least phenotypic variance 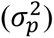 was recorded for HI (0.0) whereas LA recorded the topmost values of 8178.9 (NS), 8175.8 (PrADS) and 8090.1 (PoADS) (Table 5). Generally, the phenotypic variance appeared to be higher than the genotypic variance for all the traits across the three different conditions for all the genotypes (Table 5).

**Table 5:** Estimation of genetic parameters in eleven traits of 225 sorghum genotypes grown in multiple locations.

| Traits | $\sigma_e^2$ | | | $\sigma_g^2$ | | | $\sigma_{ge}^2$ | | | $\sigma_p^2$ | | |
| --- | --- | --- | --- | --- | --- | --- | --- | --- | --- | --- | --- | --- |
|  | NS | PrADS | PoADS | NS | PrADS | PoADS | NS | PrADS | PoADS | NS | PrADS | PoADS |
| Days to anthesis | 15.6 | 10.3 | 13.1 | 12.6 | 20.5 | 15.9 | -2.8 | -2.4 | -1.7 | 25.4 | 28.5 | 27.3 |
| Days to maturity | 36.5 | 46.3 | 42.6 | 43.6 | 62.7 | 57.6 | 2.3 | -9.5 | -6.2 | 82.4 | 99.4 | 94.0 |
| Plant height (cm) | 327.3 | 583.0 | 1893.3 | 1310.8 | 1259.9 | 859.6 | -62.9 | -51.9 | 478.8 | 1575.1 | 1791.0 | 3231.7 |
| Leaf area (cm<sup>2</sup>) | 2324.2 | 4365.2 | 4096.5 | 6327.0 | 3188.4 | 1819.3 | -472.3 | 622.2 | 2174.3 | 8178.9 | 8175.8 | 8090.1 |
| Chlorophyll content (mgcm<sup>-2</sup>) | 35.2 | 30.0 | 50.0 | 147.4 | 28.8 | 24.4 | 8.7 | -1.5 | -7.0 | 191.4 | 57.3 | 67.4 |
| Stomatal conductance (mmol m<sup>-2</sup> s<sup>-1</sup>) | 30.7 | 33.9 | 1179.8 | 2028.9 | 1834.9 | 1336.4 | -6.7 | 61.8 | 18.5 | 2052.9 | 1930.6 | 2534.6 |
| Stay green | 0.2 | 0.3 | 0.4 | 0.4 | 0.5 | 0.3 | -0.1 | -0.1 | 0.0 | 0.6 | 0.7 | 0.6 |
| 1000- seed weight (g) | 35.8 | 76.7 | 70.3 | 27.9 | 17.2 | 18.6 | 7.6 | 6.0 | -0.2 | 71.2 | 99.9 | 88.7 |
| Biomass yield (t/ha) | 9.7 | 6.4 | 8.9 | 0.3 | 0.4 | 0.5 | 0.1 | -0.1 | 2.3 | 10.1 | 6.7 | 11.7 |
| Harvest Index | 0.0 | 0.0 | 0.0 | 0.0 | 0.0 | 0.0 | 0.0 | 0.0 | 0.0 | 0.0 | 0.0 | 0.0 |
| Grain weight (t/ha) | 0.2 | 0.1 | 0.1 | 0.4 | 0.1 | 0.2 | 0.1 | 0.0 | 0.0 | 0.7 | 0.2 | 0.3 |

| Traits | GCV |  |  | PCV |  |  | H<sup>2</sup> (%) |  |  | GA |  |  |
| --- | --- | --- | --- | --- | --- | --- | --- | --- | --- | --- | --- | --- |
|  | NS | PrADS | PoADS | NS | PrADS | PoADS | NS | PrADS | PoADS | NS | PrADS | PoADS |
| Days to anthesis | 5.6 | 6.6 | 6.2 | 7.9 | 7.8 | 8.1 | 49.7 | 72.1 | 58.2 | 515.7 | 792.5 | 626.8 |
| Days to maturity | 5.6 | 6.6 | 6.5 | 7.7 | 8.4 | 8.3 | 52.9 | 63.0 | 61.3 | 990.2 | 1294.5 | 1223.5 |
| Plant height (cm) | 15.9 | 16.7 | 12.5 | 17.4 | 19.9 | 24.2 | 83.2 | 70.3 | 26.6 | 6803.6 | 6132.9 | 3114.9 |
| Leaf area (cm<sup>2</sup>) | 19.9 | 17.4 | 13.1 | 22.6 | 27.8 | 27.6 | 77.4 | 39.0 | 22.5 | 14411.8 | 7264.1 | 4166.7 |
| Chlorophyll content (mgcm<sup>-2</sup>) | 26.9 | 14.1 | 11.9 | 30.6 | 19.8 | 19.8 | 77.0 | 50.2 | 36.2 | 2194.9 | 783.2 | 612.7 |
| Stomatal conductance (mmol m<sup>-2</sup> s<sup>-1</sup>) | 23.5 | 24.9 | 20.8 | 23.6 | 25.6 | 28.7 | 98.8 | 95.0 | 52.7 | 9224.4 | 8602.6 | 5468.1 |
| Stay green | 19.5 | 28.7 | 23.3 | 23.6 | 35.2 | 34.7 | 68.2 | 66.8 | 45.2 | 105.0 | 114.3 | 71.8 |
| 1000- seed weight (g) | 10.9 | 11.0 | 10.5 | 17.4 | 26.5 | 23.0 | 39.1 | 17.2 | 21.0 | 679.9 | 354.9 | 407.0 |
| Biomass yield (t/ha) | 4.9 | 10.9 | 9.1 | 29.7 | 43.6 | 42.4 | 2.7 | 6.3 | 4.6 | 17.9 | 33.6 | 32.6 |
| Harvest Index | 10.6 | 9.4 | 8.5 | 23.5 | 27.2 | 27.1 | 20.2 | 12.0 | 9.8 | 2.5 | 1.6 | 1.4 |
| Grain weight (t/ha) | 17.7 | 15.2 | 18.1 | 22.9 | 24.0 | 22.3 | 59.6 | 40.2 | 65.3 | 104.1 | 34.8 | 77.6 |

$\sigma_e^2$ , $\sigma_g^2$ , $\sigma_p^2$ = environmental, genotypic, and phenotypic variance, PCV = phenotypic coefficient of variation, GCV = Env. = environment; genotypic coefficient of variation; $H_b^2$ = broad-sense heritability; GA = Genetic advance; NS, PrADS, PoADS see above

The PCV and GCV for all the genotypes across environments were divided into three categories (above 20% was high, 10–20% was medium, and below 10% was low). Under NS condition, the GCV ranged from 4.9 for BM to 26.9 for SPAD. In addition, SPAD (26.9) and SC (23.5) recorded high GCV values, whereas medium values were recorded by LA (19.9), SG (19.5), GY (17.7), PH (15.9), TKW (10.9) and HI (10.6). Low GCV values were recorded for BM (4.9), DM (5.6) and DA (5.6) under NS condition. Under PrADS, the GCV values ranged from 6.6 for DM to 28.7 for SG. The following traits recorded high values for GCV including SG (28.7), and SC (24.9), while others recorded moderate GCV values including LA (17.4), PH (16.7) and GY (15.2), except DM (6.6), DA (6.6) and HI (9.4) which recorded low GCV values under PrADS condition. Under PoADS, GCV values ranged from 6.2 for DM to 23.3 for SG. High GCV values were recorded for SG (23.3), and SC (20.8) while the following recorded moderate GCV values GY (18.1), LA (13.1), PH (12.5), SPAD (11.9) and TKW (10.5). All the rest of the traits recorded low GCV value (Table 5).

The PCV values ranged from 7.7 for DM to 30.6 for SPAD. All traits recorded high PCV values including SPAD (30.6) and BM (29.7) except DA (7.9) and DM (7.7) which recorded low PCV values under NS conditions. Under PrADS conditions, PCV values ranged from 7.8 for DA to 43.6 for BM. All traits recorded high PCV values under PrADS conditions except DA (8.4) and DA (7.8). Under PoADS condition, PCV values ranged from 8.1 (DA) to 42.4 for BM. All traits recorded high PCV values under PoADS condition except DM (8.3) and DA (8.1) which recorded low PCV values (Table 5).

### Heritability and Genetic Advance

The estimation of broad sense heritability 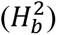 offers a descriptive measure that can be used to assess the usefulness and precision of results from cultivar evaluation trials. In the present study, under NS condition, the 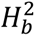 ranged from 2.7% for BM to 98.8% for SC. High values were recorded for SC (98.8%), PH (83.2%), LA (77.4%), SPAD (77.0%), and SC (68.2%). Moderate values were recorded for GY (59.9%), DM (52,9%), DA (49.7%) and TKW (39.1%) whereas low values were recorded for HI (20.2%) and BM (2.7%) under NS condition. Under the PrADS condition, the 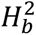 values from 6.3% for BM to 95.0% for SC. High values were recorded for most traits including SC (95.0%), and DA (72.1%). Moderate 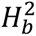 values were recorded for SPAD (50.2%), GY (40.2%) and LA (39.0%). In contrast, low values were recorded for TKW (17.2%), HI (12.0%), and BM (6.3%) under PrADS condition. Under PoADS conditions, the 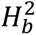 values ranged from 4.6% for BM to 65.3% for GY. High 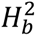 values were recorded for two traits including GY (65.3%), and DM (61.3%), whereas moderate values were recorded most traits including DA (58.2%), SC (52.7%), SG (45.2%), SPAD (36.2%), and low values were recorded for PH (26.6%), LA (22.5%) and TKW (21.0%), HI (9.8%) and BM (4.6%) (Table 5).

Under NS condition, GA values ranged from 2.5 for HI to 14411.8 for LA. Highest values were recorded for LA (14411.8), and SC (9224.4) whereas the lowest values were recorded for HI (2.5) and BM (17.9). Under PrADS condition, GA values ranged from 1.6 for HI to 8602.6 for SC. Highest values were recorded for SC (8602.6) and LA (7264.7) whereas lowest values were recorded for HI (1.6) and BM (33.6). Under PoADS condition, GA values ranged from 1.4 for HI to 5468.1 for SC. Highest values were recorded for SC (5468.1) and LA (4166.7) whereas lowest values were recorded for HI (1.4) and BM (32.6) (Table 5). High and moderate heritability values coupled with high genetic advances were found in almost all the traits, except BM and HI (Table 5).

### Pearson’s correlation

To evaluate the relationships between assessed parameters (traits), Pearson correlations were performed across drought treatments for both locations and presented in Figure 2. Under NS conditions, GY showed strong positive correlations to BM (r = 0.44; p < 0.001) and HI (r = 0.88; p < 0.001) across environments (Figure 1A). DM (r = 0.26; p < 0.001), LA (r = 0.24; p < 0.001), SG (r = 0.19; p < 0.001) and BM (r = 0.24; p < 0.001) were positively related to DA. In addition, DM was positively and significantly correlated with PH (r = 0.37; p < 0.001) (Figure 2A).

**Figure 2:**
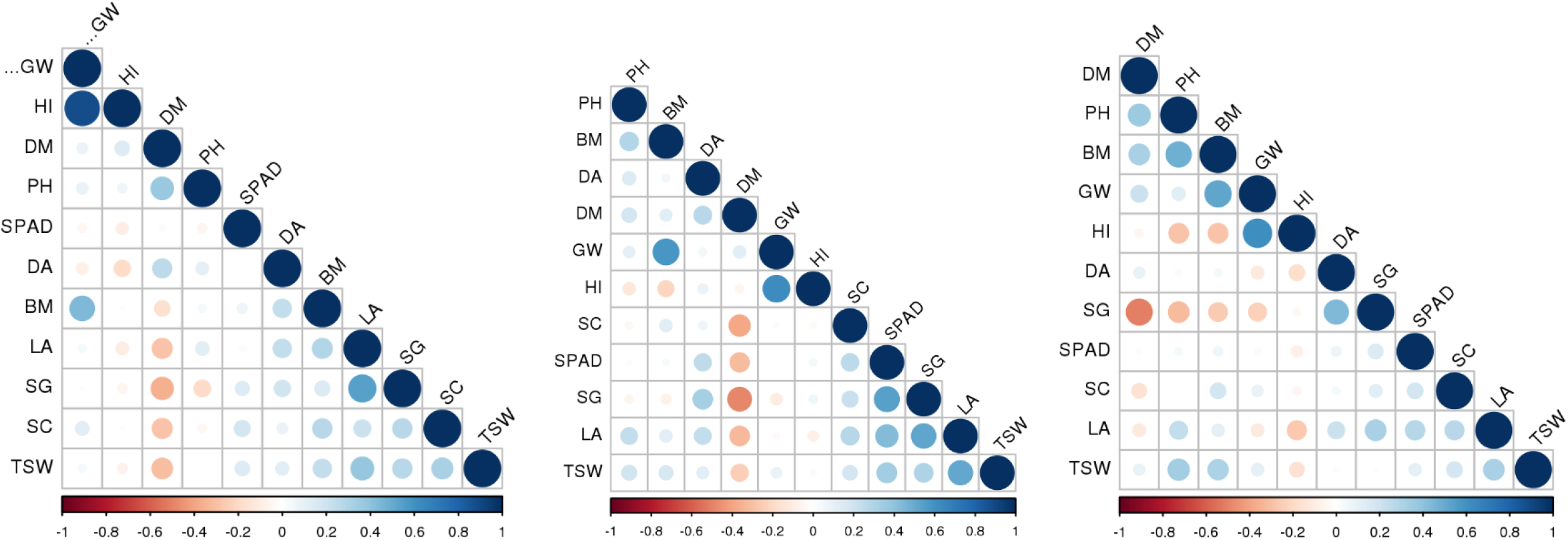
Pearson correlation coefficients (r) of grain yield and other agronomic traits among sorghum genotypes under non-stress (A), pre-anthesis (B) and post-anthesis (C) drought conditions across test environments. GW=grain yield (t/ha); DA=days to anthesis; DM=days to maturity; PH=plant height, LA=leaf Area; SPAD= chlorophyll content; SC= stomatal conductance; SG=stay green; TSW; thousand seed weight; BM=aboveground biomass and HI=harvest index.

Under PrADS, GY was positively and moderately correlated with BM (r = 0.36; P < 0.001) and HI (r = 0.62; P < 0.001) across environments (Figure 2B). DA was positively and weakly correlated with DM (r = 0.27; P < 0.001), PH (r = 0.14; P < 0.05), LA (r = 0.23; P < 0.001), SPAD (r = 0.25; P < 0.001) and SG (r = 0.32; P < 0.001). The data, DM was positively and weakly associated with PH (r = 0.19; P < 0.001) and negatively correlated with LA (r = 0.33; P < 0.001), SPAD (r = 0.32; P < 0.001), SC (r = 0.39; P < 0.001) and TKW (r = 0.25; P < 0.001). PH was positively associated with LA (r = 0.24; p < 0.001), TKW (r = 0.20; p < 0.01), and BM (r = 0.30; p < 0.001) under PrADS conditions across environments. The BM was negatively and strongly correlated with HI (r = 0.63; P < 0.001) (Figure 2B).

Under PoADS conditions, GY was positively correlated to DM (r = 0.21; P < 0.05), BM (r = 0.52; p < 0.001), HI (r = 0.61; P < 0.001) and negatively correlated to SG (r = 0.24; P < 0.001) (Figure 2C). PH was positively correlated with LA (r = 0.26; P < 0.001), BM (r = 0.47; P < 0.001), and negatively correlated to SG (r = 0.22; P < 0.001). LA was positively and weakly correlated with SPAD (r = 0.28; P < 0.001), SC (r = 0.26; P < 0.001), SG (r = 0.32; P < 0.001), TKW (r = 0.36; P < 0.001) and negatively correlated with HI (r = 0.29; P < 0.001). The BM data was negatively associated with HI (r = 0.29; P < 0.001) under PoADS conditions (Figure 2C).

## DISCUSSIONS

The environment in the current study represented an environment with water limitation starting during the vegetative growth in one environment and starting during anthesis in the other and continuing with severe drought stress to crop physiological maturity. This type of environment is representative of sorghum production systems in SSA. The differences between the two environments (normal and stressed) are reflected in the observed patterns of PrADS and PoADS grain yield production (Table 4 and Figure 1). The present study found significant variations in grain yield and all measured traits among the genetically diverse sorghum genotypes (Table 3). The significant genotype differences observed among the studied sorghum genotypes for grain and biomass yield, days to anthesis, and quality traits allowed selection of suitable dual-purpose genotypes (Table 3). Also, genotype × environment × water regime interaction effect was significant for most of the measured traits, suggesting that the observed variations in the assessed agro-morphological traits were largely due to genotypic differences, environment, and water regime (Table 3). Several studies on sorghum utilizing a variety of morphological parameters confirmed our findings about the presence of high genetic diversity in the African sorghum collections. (Motlhaodi et al., 2014; Habyarimana et al., 2004; Amelework et al., 2015). The relevant observation of high variation coefficients was corroborated by Motlhaodi et al. (2014) and Amelework et al. (2015), which further demonstrated the existence of a massive heterogeneity among the landraces observed in sorghum.

The markedly lower grain yields (Table 4), and the reduced plant heights and delayed flowering, reduced leaf area, chlorophyll content and stomatal conductance (Supplemental Table S2) observed in PrADS and PoADS test environments, relative to the NS environments in each location were of similar magnitude as those observed across a wide range of sorghum genotypes tested under varying drought conditions (Harris et al., 2007, Burke et al., 2018, Tsago et al., 2014; Emendack et al. 2018; de Souza et al., 2020). In addition, the yield levels under both PrADS and PoADS conditions were higher than mean average in farmers’ field across Africa (< 1.0 t/ha), (FAOSTAT 2020), suggesting that our results are relevant for targeting the drought prone sorghum production systems in this region.

The evaluated sorghum genotypes significantly exhibited variability for number of days to anthesis under drought-stressed condition (Table 4 and Figure 1a). Compared to non-stressed environment, sorghum genotypes in drought stressed treatments of PrADS had accumulated on average 7.4 % less days to anthesis pre-anthesis, whereas post-anthesis DA accumulation was 2.0 % less across environments (Supplementary Table 2). The reduction for days to anthesis under drought stress compared to non-stress is a form of genotypic adaptation to drought stress and according to Yohannes et al., (2015) earliness prompts the crops to flower early to escape drought stress which often leads to reduced yield or complete crop failure. According to Yohannes et al. (2015), early flowering indicates early maturity, and according to our findings, the early flowering genotypes such as IS 8268, ICNSL2014-022-8 and SSV20071012, can be selected for earliness as ways of drought escape mechanism. Days to maturity differed significantly (p ≤ 0.01) among the studied genotypes and varies from 90 to 146 days under NS, 81 to 151 under PrADS condition and 89 to 147 days Under PoADS (Table 4 and Figure 1). Similar days to maturity range was reported by Angarawai et al., (2021) and Assefa et al., (2010). The maturity duration of the sorghum depends on the cultivar and climatic situation (Angarawai et al., 2021). Sorghum genotypes from the Sahelien and Sudanian agro-ecologies were typically early to medium maturing (70-89 days to 50% flowering) and associated with short (2 m) or medium (2.5 m) plant heights due to the relatively low rains prevalent in the agro-ecology (≤ 500mm), while collections from the Northern and Southern Guinea savannah were typically late and tall, recording more than 110 days to attain days to 50% flowering and plant heights greater than 3.5 m, respectively. Although there is a negative yield penalty associated with earliness as reported by several studies, genotypes that mature early have been reported to contribute to yield stability in arid regions where terminal drought stress occurs (Assefa et al., 2010; Angarawai et al., 2021). Therefore, high grain producing genotypes with a medium height of 2 meters and a medium maturity of 100 days should be the current focus of sorghum hybrid parent development for sustainable hybrid sorghum production in drought-prone locations.

Plant heights across the environments were significantly different and this was greatly influenced by the genotype. The current study revealed positive correlations between plant height with days to anthesis and leaf area under drought stress conditions (Figure 2B and 2C) suggesting an increase in plant height likely influenced the days to flowering and enhanced leaf area. However, we observed an increase in plant height (7.0%) under pre-anthesis drought stress condition and a reduction in plant height (2.8%) under post-anthesis. According to Blum (1996) reduced plant height is a symptom of moisture stress. In this study, the sorghum genotypes that are able to escape drought (days to anthesis ≤ 60 days) are usually short in stature such as AS 71 (77.0cm), P9402 (85.6cm), GP11BR (93.9cm) and E 29 (94.3cm). These genotypes have some sort of genotypic adaptation to enable them to utilize the little moisture available to complete their life cycle without compromising yields. Although the current study found no significant relationship between plant height and grain yield, other studies have found a strong relationship between plant height and grain yield (Kapanigowda et a., 2013). Tall plants were linked to longer maturation times and higher yields under ideal conditions, but they were linked to lower yields under sub-optimal and unfavorable growing conditions (Angarawai et al., 2021). Consequently, the phenotypically varied plant heights displayed by the examined sorghum genotypes under drought stress conditions certainly influenced their water consumption and the severity of water deficit encountered.

The rate of transpiration and gas exchange through leaf stomata is approximately determined by leaf stomatal conductance (Vico et al., 2013). Plants use a reduction in leaf stomatal conductance as a defense mechanism to escape or tolerate drought (Lawlor, 2013). As water regimes varied among the locations, different stomatal conductance patterns were seen. In this research, control treatments of sorghum genotypes achieved higher stomatal conductance than drought stressed genotypes. The stomata on upper leaves of drought-stressed genotypes were more open in PoADS condition (µ = 175.68 mmol m-^2^ s^-1^) than in PrADS plants (µ = 171.92 mmol m-^2^ s^-1^) (Table S2 and Figure 1g). It is suggested that the plants’ stomata remained open under PoADS condition even after bulk leaf turgor was lost. From these results and from relationships between leaf stomatal conductance and stage of panicle development, it is concluded that the tendency of stomata to remain open despite water stress is related to the presence of an emerged panicle (Vico et al., 2013). Similar pattern was observed for leaf chlorophyll content of studied sorghum genotypes and were influenced by the water treatment which was more pronounced in PrADS condition (µ = 38.2 mgcm^-2^) when compared to PoADS condition (µ = 41.4 mgcm^-2^) (Table S2 and Figure 1f). El Sabagh et al., (2017) reported similar striking genotypic differences of stomatal conductance and leaf chlorophyll content.

In addition, the inverse relationship between leaf chlorophyll content with days to maturity suggested that leaf chlorophyll content index decreases in late maturing genotypes with extended growth phase. The high positive significant correlation coefficients observed under PoADS condition between leaf area and physiological traits (leaf stomatal conductance and chlorophyll content) suggested that the effect of moisture stress on the leaf is related to photosynthetic capability and stomatal conductance of the genotypes. According to Wang et al., (2016), the relationship between stomatal conductance and a plant’s photosynthetic capacity, can be assessed by measuring the chlorophyll content of the leaves. In the current study, a correlation between stomatal conductance and chlorophyll content was not evident so that none of the drought stress treatments could be identified as having any positive influence on the two processes simultaneously. Despite showing superior performance under PrADS and PoADS conditions for leaf stomatal conductance and chlorophyll content, the breeding lines from ACCI-SA (AS 71 and E 119) and IAR-NG (SAMSORG 7), did not have the highest yields. Sorghum plant response to water stress has been linked to decreasing water loss by controlling stomatal activities under drought conditions, allowing the plants to prevent dehydration and physiological decline (Ncama et al., 2022). Decreased stomatal conductance and reduced photosynthetic activity are indicators of effective drought avoidance in the studied sorghum collection.

Stay green is an important drought-adaptation trait that allows plants to retain their leaves in an active photosynthetic state when subjected to soil moisture shortage (Rosenow et al., 1996). According to Mahalashmi and Bidinger (2002), stay-green genotypes of sorghum retain more green leaf area and higher levels of stem carbohydrates while also being less prone to lodging and more resistant to charcoal rot. In the current study, performance under drought stress conditions (PrADS and PoADS) was significantly associated with green leaf area and days to maturity. Higher heritability estimates for the examined stay-green trait provide compelling evidence that the observed phenotypic differences were mostly influenced by genetic factors. Differences among genotypes for stay green trait was significantly affected by the water regime (Table 3). Drought stress under pre- (µ = 2.4) and post-anthesis (µ = 2.2) growth stages caused leaf senescence which results in degradation of chlorophyll and disorganization of the photosynthetic apparatus. From the results, the genotypes did not show superior yield performances for stay green especially under post-anthesis drought stress. From the results, is appears that the stay-green trait independent of grain yield. According to Xu et al., (2000), where drought is not effectively expressed, particularly due to timing and degree of moisture stress, and the existence of significant genotype environment interactions, field measurement of the stay green trait can be challenging and unreliable.

In our assessment, significant difference (p ≤ 0:01) was found for thousand seed weight which varied from 25 g (Juar) to 75 g for control plants and between 18 to 60.8 g for pre-anthesis drought stressed plants, and between 20.8 g to 58.6 g for post-anthesis drought stressed plants (Table 4 and Figure 1i). According to several research, seed weight is an essential tool for evaluating morphophysiological features that are connected to yield (Motlhaodi et al., 2014; Angarawai et al., 2021). The low coefficient of variation value (18.4%) suggested that this trait may be a reliable selection criterion for drought resistance. The considerable decline in yield-related parameters, such as 1000-seed weight, observed under pre- and post-anthesis drought stress indicated that grain yield was impacted by drought stress at these periods. The decrease in 1000-seed weight due to drought stress in current study was due to shriveling of the seed.

Large biomass production coupled with high harvest index is likely to result in efficient grain production. As pointed out by Angarawai et al., (2021), the local landraces of sorghum are tall, late maturing, locally adapted, and exhibit low harvest index (about 30%). High yielding cultivars with efficient partitioning of dry matter into the components is an important attribute of modern improved sorghum cultivars. The results of the current study, Due to the effective drought avoidance mechanisms in most of the studied genotypes, the crop can withstand the severe drought conditions that can occur in arid and semi-arid locations, such as the dry parts of West Africa, where the majority of the genotypes were collected. The highest yielding genotypes with dual-purpose features were landraces under pre-anthesis drought stress conditions with the following grain and above-ground biomass yield respectively: CSRO1 (2.6 and 7.3 t/ha), TUN BUMAN MAIDUGURI (2.8 and 5.8 t/ha), ICNSL2014-021-1 (2.7 and 7.4 t/ha), S7-Lata/RIB/BC1-3-1-1-V (2.3 and 10.8 t/ha) and JAR KAURA (2.4 and 9.4 t/ha). Under the post-anthesis drought stress, the genotypes with highest performance for both grain and above-ground biomass yield include GAGARAU - 4 (4.2 and 12.2 t/ha), KAURA SHORT PANICLE-1 (4.1 and 12.3 t/ha), KAURA MAI BAKI KONA (3.9 and 11.4 t/ha) and YAR LAZAU (3.8 and 14.5 t/ha). The positive significant correlation coefficients between grain yield with above-ground biomass suggests that the two traits can be simultaneously improved which is consistent with earlier reports in sorghum by Makanda (2017) and Alam et al. (2001). The PCV was higher than the GCV for both traits, and higher heritability combined with high genetic advance revealed the presence of additive genetic control for both traits.

The harvest index measures the physiological efficiency of the plants. Breeders have recognized the importance of a favourable harvest index in terms of partitioning of photosynthate to economically important plant part. Harvest index in local and hybrids sorghum reported to be 29 and 50 per cent, respectively (Piper and Kulakow 1994). Thus, there is a greater need for genetic manipulation to increase the harvest index. Productivity of sorghum can be improved by enhancing the biological yield without losing the harvest index. The harvest index of high-performance genotypes such as CSRO1 (0.30), TUN BUMAN MAIDUGURI (0.33) under pre-anthesis drought stress and GAGARAU - 4 (0.27), KAURA SHORT PANICLE-1 (0.28), under post-anthesis drought stress revealed the presence of significant variation for harvest index under drought stress. This information underscores the need for enhancing the overall genetics of the identified high performing and drought-tolerant landraces in order to improve their drought adaptive traits. This will need to be accomplished either by choosing the relevant landraces under intense cultivation and enhancing their agronomic qualities or by introgressing the distinctive traits into farmer-preferred cultivars.

### Genetic Parameter Analysis

Genotypic variance is a metric that gauges the strength of genetic and heritable variation (Burton et al., 1953). Given the significant genetic diversity in this group, selection can be successfully used. The phenotypic variance values for all of the characters were higher than the corresponding genotypic variance values, indicating that most of the trait’s expression is impacted by the environment. These findings of our research were supported by previous reporters (Sami et al., 2013; Yohannes et al., 2015; Hamidou et al., 2018). The genotypic coefficient of variation (GCV) and phenotypic coefficient of variation (PCV) were medium to high for most of the measured traits. In general, the PCV was higher than the GCV for all measured traits. This suggested that selection will be fruitful for the development of traits.

The broad-sense heritability is the proportion of phenotypic variance that is attributable to an overall genetic variance for the genotype (Schmidt et al., 2019). In this current research, almost all the traits related to yield recorded medium to high broad-sense heritability values except harvest index, and above-ground biomass yield that recorded low heritability. Under Pre-anthesis drought stress, traits such as days to anthesis, plant height, stomatal conductance, and stay-green recorded higher heritability estimates than grain yield and can be used in indirect selection in breeding for improved pre-anthesis drought resistance and can help optimize grain yield productivity in stress environments. In contrast, traits such as days to maturity and grain yield recorded high heritability estimates under post-anthesis drought stress environments. According to Alam et al., (2022) the estimation of heritability of the tested genotypes paired with high genetic advance (GA) is more accurate and effective for the selection of desirable traits for a group of the population. Hence, those traits with high values for GCV, PCV, heritability, and genetic advance imply the presence of many genes with additive effects that enable efficient selection for improving traits directly.

### Association Analysis

According to Hallauer et al., (2010), a plant breeder must give the utmost consideration to the interaction of different yield components when choosing selection criteria. Under all conditions, our findings revealed the presence of positive and significant correlation of days to anthesis with days to maturity and plant height indicates that these traits can be directly selected for improvement. The chlorophyll content was positively associated with stay green, stomatal conductance and under all conditions. However, under pre-anthesis drought stress, a positive relationship was observed with days to anthesis, and a negative correlation was observed with days to maturity. The findings revealed that photosynthesis was at first undisturbed at the onset of water stress before flowering which becomes critical due to prolonged exposure to drought stress. According to Assefa et al., (2010) and Wang et al., (2016), drought stress disrupts the basic organizational structure, which prevents carbon uptake and harms the photosynthetic apparatus. In addition, the increasing frequency and intensity of drought stress in pre- and post-flowering conditions is likely the reason contributing to the weak and negative correlation between stay-green and grain yield. Thousand seed weight numbers detected positive and moderate highly significant correlation with grain yield across all irrigation regimes. However, a negatively strong correlation was noted with the days to maturity under non-stressed and pre-anthesis drought condition and weak positive association under post-anthesis drought stress condition. The variable association observed under pre- and post-anthesis drought stress may suggest that the consideration have to be given when selecting for these traits under different water regimes. Significant positive correlation was observed between above-ground biomass and grain yield. Harvest index and grain yield were also observed to be positively associated. In contrast, above-ground biomass recorded negative correlation with harvest index under both pre- and post-anthesis drought stress. Based on these observations, it is suggested that sorghum may be improved by using a selection criterion that took biomass and harvest index into account.

## Conclusion

The sorghum genotypes differed significantly for most of the parameters collected in this study. Due to the effective drought avoidance mechanisms in most of the studied genotypes, the crop can withstand the severe drought conditions that can occur in arid and semi-arid locations, such as the dry parts of West Africa, where the majority of the genotypes were collected. The genotypes CSRO1 (2.93 t/ha), Tun Buman Maiduguri (2.80 t/ha) and ICNSL2014-021-1 (2.69 t/ha) showed good resistance to pre-anthesis drought stress, and was desirable for earliness, maintenance of photosynthetic capacity and stomatal conductance under pre-anthesis drought stress. Under post-anthesis drought stress conditions: Danyar Bana (4.21 t/ha), GAGARAU - 4 (4.15 t/ha) and Kaura Short Panicle-1 (4.08 t/ha) revealed good drought resistance. The identified genotypes with superior yield performance represent really valuable material that can be deployed for cultivation in the dry and semiarid ecologies.

## Acknowledgement

The authors sincerely acknowledge the Directorate of Research and Innovation (DRI), Ahmadu Bello University, Zaria, for providing financial support for this research through the TETFund Institution-Based Research Grant. Appreciation is also extended to the Institute for Agricultural Research (IAR), Samaru for providing the research facilities and conducive environment necessary for the successful execution of the study. The technical and institutional support received from Ahmadu Bello University in facilitating the implementation of the research project is also gratefully acknowledged.

